# DINAVID: A Distributed and Networked Image Analysis System for Volumetric Image Data

**DOI:** 10.1101/2022.05.11.491511

**Authors:** Shuo Han, Alain Chen, Soonam Lee, Chichen Fu, Changye Yang, Liming Wu, Seth Winfree, Tarek M. El-Achkar, Kenneth W. Dunn, Paul Salama, Edward J. Delp

## Abstract

**Background:** The advancement of high content optical microscopy has enabled the acquisition of very large 3D image datasets. Image analysis tools and three dimensional visualization are critical for analyzing and interpreting 3D image volumes. The analysis of these volumes require more computational resources than a biologist may have access to in typical desktop or laptop computers. This is especially true if machine learning tools are being used for image analysis. With the increased amount of data analysis and computational complexity, there is a need for a more accessible, easy-to-use, and efficient network-based/cloud-based 3D image processing system.

**Results:** The Distributed and Networked Analysis of Volumetric Image Data (DINAVID) system was developed to enable remote analysis of 3D microscopy images for biologists. DINAVID is a server/cloud-based system with a simple web interface that allows biologists to upload 3D volumes for analysis and visualization. DINAVID is designed using open source tools and has two main sub-systems, a computational system for 3D microscopy image processing and analysis as well as a 3D visualization system.

**Conclusions:** In this paper, we will present an overview of the DINAVID system and compare it to other tools currently available for microscopy image analysis.

## Introduction

Recent progress in microscopy technology has enabled the acquisition of large 3D volumetric data [1, 2], including 3D multi-spectral data, using fluorescence imaging. The analysis of such data presents several challenges. The first major challenge is the accurate quantification of features in objects of interest. For example, image analysis tools are commonly used to quantify the amount of fluorescence of different protein stains within each cell [3]. CellProfiler [4] and ImageJ [5], which we will discuss in more detail below, are examples of such popular image analysis packages frequently utilized in the biology community. This first challenge is exacerbated by the characteristics of biological structures, that include crowding of structures, blurred boundaries, and various noise types. Nonetheless, the tremendous expansion and versatility of machine learning and deep learning has allowed for the development of more powerful tools that are capable of extracting and quantifying important biological information such as the presence of important proteins. For example, U-Net [6] is a deep learning architecture developed originally for segmentation of biomedical data and is very commonly used for segmentation of microscopy images. In fact, both ImageJ [5] and CellProfiler [4] have been making progress in incorporating U-Net [6] and other machine learning tools. In some cases manual installation of libraries is needed which limits use for non-expert users. In other cases, a user needs to provide a set of original microscopy images and corresponding groundtruth images to train the network.

The second challenge is to obtain representative and effective microscopic visualization and imaging of biological structures which are vital for the analysis and understanding of related biological processes [7]. Traditional 2D visualization often lacks important perspectives, and thus effective 3D visualization is needed for a more complete understanding of the data. A variety of ways have been used to achieve 3D visualization of an image volume, such as maximum intensity projection (MIP), three-dimension views, or cross-sectional views [8]. We will review the commonly used visualization tools in the next section.

A third challenge is the computational resources needed for large 3D multi-channel image analysis. 3D microscopy volumes often range in size from several gigabytes (GBs) to terabytes (TBs) of data, and consequently require more computational resources (e.g. many GPUs) than a biologist may have access to in typical desktop or laptop computers, especially if machine learning tools are being used for image analysis. With this increased amount of data analysis and computational complexity, there is a need for a more accessible, easy-to-use, and efficient network-based/cloud-based 3D image visualization and processing system. Another factor limiting the analysis of image volumes is friendly software. This can include unfriendly graphic user interfaces. The need to install multiple software packages, sometimes from a command line, can also be a burden to the user experience.

The Distributed and Networked Analysis of Volumetric Image Data (DINAVID) system was developed with these objectives in mind, namely to enable server/cloud-based analysis of microscopy images for biologists. The goal is to provide a user-friendly environment with a simple web interface system that biologists can use without worrying about managing the computational resources.

DINAVID is designed to support interactive visualization and exploration of large image volumes. DINAVID also supports quantitative analysis through a workflow of image pre-processing, segmentation using a variety of different techniques, quantification of the fluorescence of individual cells, and interactive data analysis. We will present an overview of DINAVID and compare it to other tools currently available for microscopy image analysis below.

## Review of Existing Systems

We overview below some of the existing systems and tools that are available for a user to analyze volumes. Similar comparisons have been made in a review article [26] for freely available software tools for single cell analysis. We will differentiate the various systems not only by their capabilities to analyze 3D volumes but also what biologists would need to manage using the systems tools with emphasis on whether the tools are network-based or require download and installation.

### “Local” Image Analysis Systems

A “local” system is a set of tools that a user would need to install on their local computer to use. Open source local image processing packages are preferred by many biologists. Some tools only support 2D visualization of 3D volumes. By 2D visualization, we mean that 3D data can be viewed as sequential slices in 2D as orthogonal planes. Examples of these are CellProfiler [4, 19], the Volumetric Tissue Exploration and Analysis tool (VTEA) [3], and Cellpose [20]. Since cross-sectional viewing could not display the objects of interest in 3D, a user needs to observe back and forth along the cross-sections to estimate the 3D surface. Thus, true interactive 3D rendering is preferred.

Examples of free-to-use software that supports interactive 3D rendering include Voxx [27], Agave [18], and Open Graphics Library (OpenGL) [28]-based systems such as ImageVis3D [17]. Imaris [21] is a commonly used commercial software tool used for microscopy image analysis and visualization. A comparison of local-based systems is summarized in Table 1. ImageJ [5] does not support 3D visualization natively but provides support for plugins for additional functionality, examples of which include the 3D Viewer plugin [29] and Big Data Viewer [24]. Other available tools that do support 3D visualization include ClearVolume [22], Napari [23], and ilastik [25].

**Table 1.**
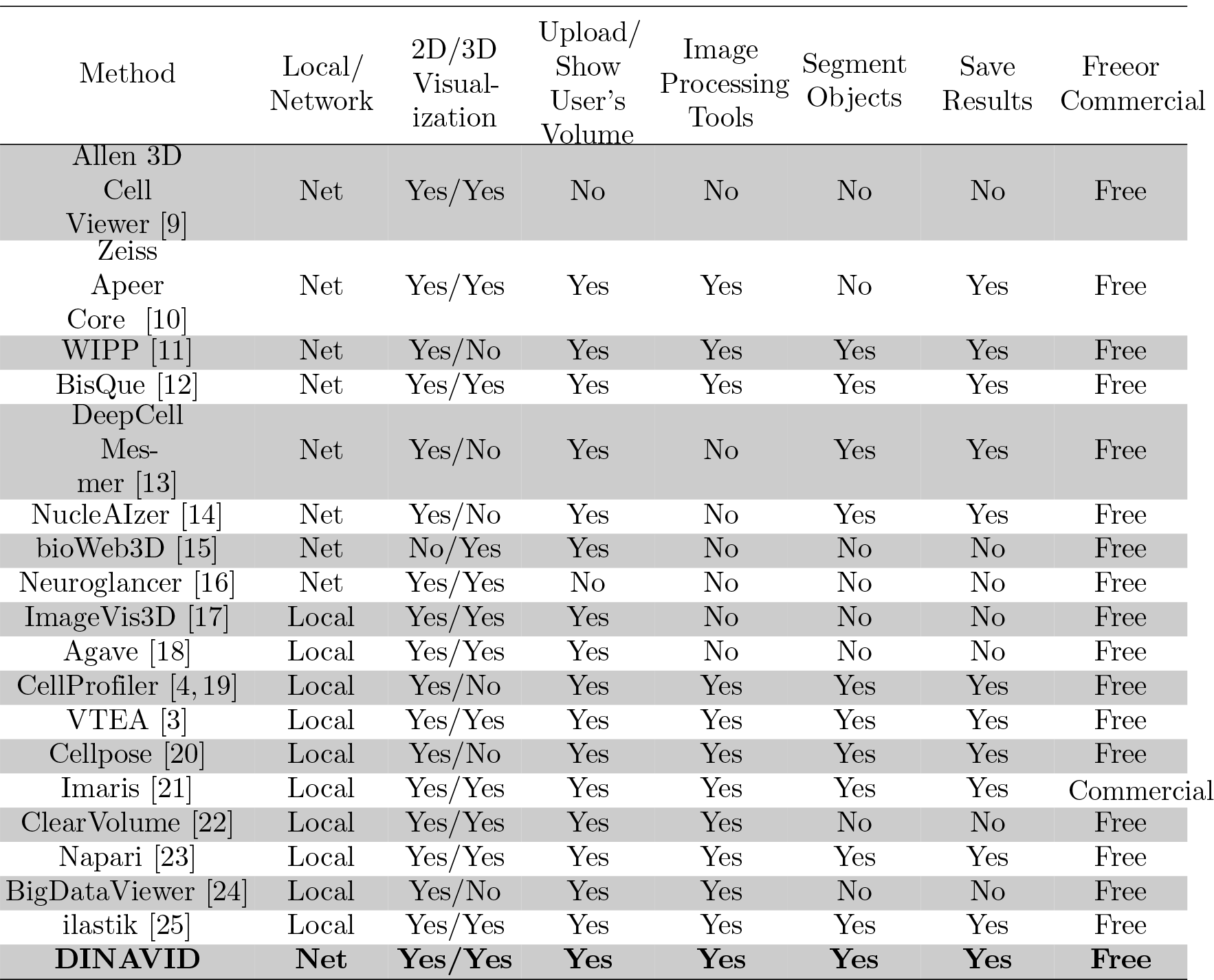
Comparison of Microscopy Image Analysis Tools

These systems all require that the software be downloaded, installed, and maintained on a local machine.

### “Network-Based” Image Analysis Systems

In contrast to “local” systems, by a network-based (or cloud-based) system we mean a system that allows a user to process and visualize 3D volumes remotely. This does not require that they install anything on their local computational resources. Users access the system via a web interface. Examples of network-based systems include Apeer [10] from Zeiss, BisQue [12], DeepCell Mesmer [13], NucleAIzer [14], 3D Cell Viewer [9] by the Allen Cell Institute, BioWeb3D [15], and Neuroglancer [16]. In addition, for the purpose of facilitating the analysis of large size image data, the Web Image Processing Pipeline (WIPP) [11] was developed by the National Institute of Standards and Technology (NIST). It was recently reported that plugins for cloud-based microscopy image analysis [30] have been developed for WIPP. A comparison of network-based systems, along with the local systems, can be found in Table 1.

Overall, most network-based solutions lack the capability of image analysis while many local-based solutions are not as versatile in their visualization functions. DINAVID bridges the gap between these two solutions by providing both intuitive visualization tools and several image processing, segmentation, and quantitative analysis capabilities.

## Architecture/Functionalities of DINAVID

The basic components of the DINAVID system are shown in Figure 1, which includes volume uploading, visualization, pre-processing, segmentation, and quantification. We will discuss each of the components in this section.

**Figure 1.**
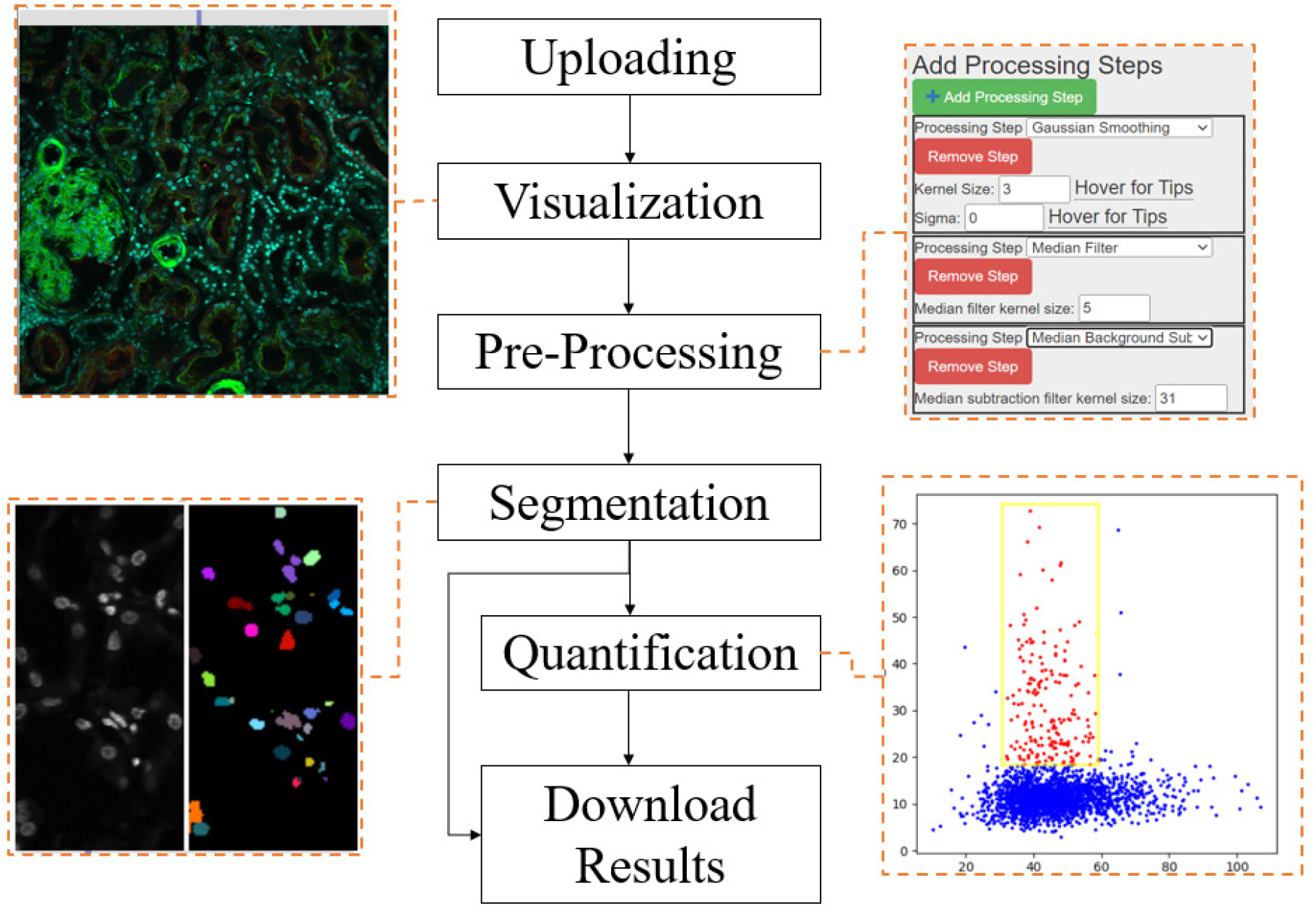
Block diagram of the DINAVID system

### File Uploading and Downloading

DINAVID supports the upload of a single 3D composite .tiff volume. The maximum number of channels supported is 19. Currently DINAVID supports volumes of up to 200GB. A user may also download their segmentation and quantification results. Upload/download speed is governed by the user’s network connection. The currently deployed version of DINAVID is connected to Internet2 [31].

### 2D and 3D Visualization

DINAVID supports both 2D and 3D visualization. As noted above by “2D visualization of 3D volumes”, we mean that a 3D volume can be viewed as sequential slices in 2D as orthogonal planes, and by 3D visualization we mean true interactive volume rendering. In the case of 2D visualization DINAVID displays a slice from a 3D section of the selected channels. Each channel in a 2D slice is assigned a default color. Alternatively, each user can assign a color to each channel individually from a provided color table. The maximum possible intensity in each channel is assigned to the selected color, while lower intensity values are assigned to colors that are scaled according to the ratio of their relative intensity values to the maximum possible intensity. A maximum projection of each channel is then used to generate the final color image that is displayed.

Users are able to scroll through single slices of a 3D volume. Users are also able to select a rectangular subregion to view the selected region at a higher magnification.

DINAVID also has the capability of interactively adjusting the gamma, brightness, and offset for each of the individual channels, defined by:

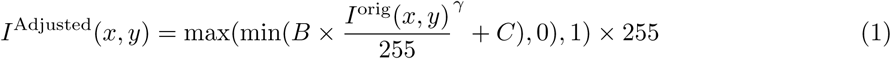

where *B, C*, and *γ* denote the brightness, offset and gamma, respectively, and *I*^orig^(*x, y*) is the intensity of the image before adjusting *B, C*, and *γ*.

For 3D visualization, DINAVID is able to render the subregion, selectable by the user, as a 3D alpha-blended volume rendering. To facilitate interactive rendering, the size of the subregion is limited, and users must select 3 of the available channels to render as the 3 channels of an RGB volume. In addition to the brightness, offset, and gamma controls as described above, the user is also able to adjust the density of the volume rendering.

Finally, users can hide and show the panels displaying parameter settings to maximize the visualization region of a microscopy volume.

### Image Pre-Processing

DINAVID supports many pre-processing operations. A user can select any combination of the operations or repeat them as needed. The main goal in this step is to reduce noise and to eliminate background effects. For a detailed description of the available pre-processing steps, please refer to Appendix B.

### Segmentation

Since cell boundaries are typically poorly defined in tissues, one approach is to segment nuclei rather than cells [3]. Biologists then characterize and classify cells on the basis of the fluorescence in the user-defined regions surrounding the nuclei [32]. DINAVID adopts this approach for segmentation. The nuclei segmentation tools currently available are 3D watershed [33], 2D watershed with joining [3], and a deep learning-based method known as DeepSynth [7]. Currently, five pre-trained versions (five inference models) of DeepSynth [7] are available for users in DINAVID.

Since some pre-processing operations and segmentation require long execution times, working directly on an entire image volume is often impractical or time consuming, especially for large volumes. A better practice is to first test the analysis on a sub-volume that shows results quickly before proceeding to process an entire volume. DINAVID is equipped with a previewing function for the pre-processing and segmentation steps that permits the previewing of the image processing or segmentation steps on user selected nuclei channels and regions of interest. This allows for users to rapidly test different image processing and segmentation settings.

### Quantification

As described previously, nuclei rather than cells are segmented as the cell boundaries are typically poorly defined in tissues [3]. DINAVID can examine the voxels from different channels in a user-defined region surrounding each nucleus and then estimates and visualizes multiple statistics. We define “the region around the nucleus” of each nucleus as the voxels that are at a user-defined distance away from the boundary of the nucleus. Using the segmented nuclei masks, voxels surrounding each nucleus are extracted using morphological dilation. Nuclei masks of each cell are dilated by a user-defined size. The difference between the resulting dilated mask and the original nuclei mask represents the region around the nucleus. The statistics include minimum intensity, maximum intensity, mean intensity, standard deviation of intensity, sum of intensity, and number of voxels, for each channel in the user-defined region. These statistics are entered into a downloadable spreadsheet for external analysis or can be displayed as 2D scatter plots where each point on the scatter plot represents a single nucleus. The axes of the scatter plot can be set to any two of the six measured statistics of any channel.

DINAVID allows users to draw a rectangular region of interest (ROI) in the image volume, the nuclei within this ROI are then displayed in the scatter plot. Alternatively, users can also draw a rectangular ROI in the scatter plot, and the relevant nuclei will be highlighted in the original image volume. This permits users to be able to determine the quantity and location of a specific type of cell inside the imaged biostructure [3, 34], based on the six statistics.

## Using DINAVID

Users interact with DINAVID via a web interface and are required to log in using the credentials that are supplied to them upon request. The user interface is easy to use and understand and is customizable. A user does not need to download and maintain components of DINAVID since all processing and data handling is done on the server/cloud. Users can upload an image volume for processing and visualization. Figure 2 shows an example DINAVID interface for selecting the parameters for 2D visualization. After the user chooses the desired channel with corresponding color to visualize a volume in Figure 2(a), the changes in brightness, gamma, and offset to the images will be reflected in shown Figure 2(b). Figure 2(c) depicts the menu of processing steps that the user may choose to apply to the uploaded image. The number of steps and the sequence of processing is customizable. Figure 2(d) shows an example of a slice where the background was subtracted using median filtering. Figure 3 shows an example interface for 3D rendered visualization with a GUI for parameter tuning appearing on the right. The user can adjust the rendered volume and its appearance interactively in real-time. An example quantitative analysis after nuclei segmentation is shown in Figure 4. Using the segmentation result obtained from watershed [33], a scatter plot is generated by specifying the statistics, which can be minimum intensity, maximum intensity, mean intensity, standard deviation of intensity, sum of intensity, or number of voxels, to plot on the vertical and horizontal axes. The corresponding nuclei from the selected region of interest are then overlaid on the original image.

**Figure 2.**
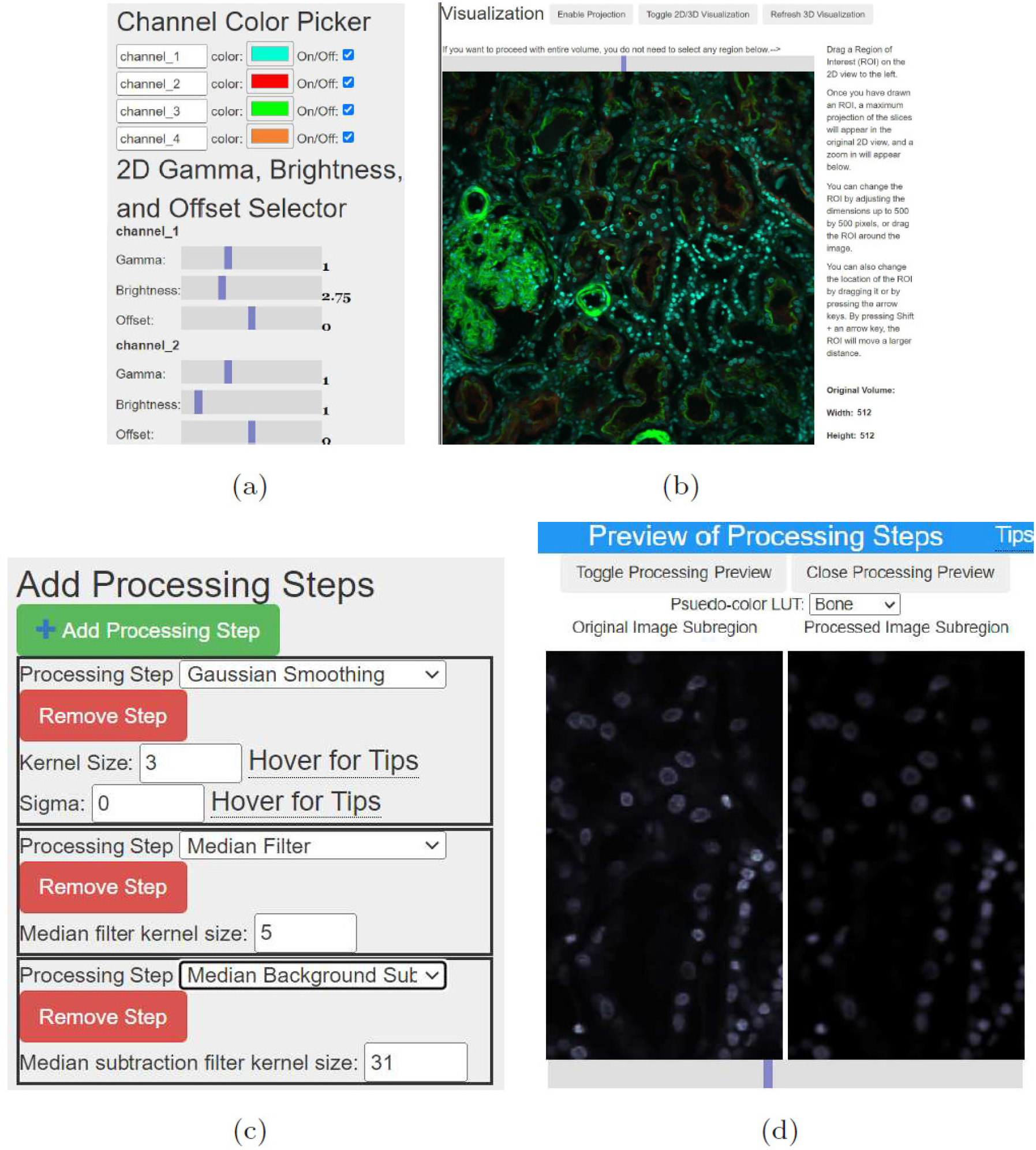
Examples of 2D image processing and visualization in DINAVID (a) Panel to adjust visualization parameters, (b) 2D Image visualization, (c) Panel to select image processing steps and adjust parameters, (d) An example of a slice after a median background subtraction operation

**Figure 3.**
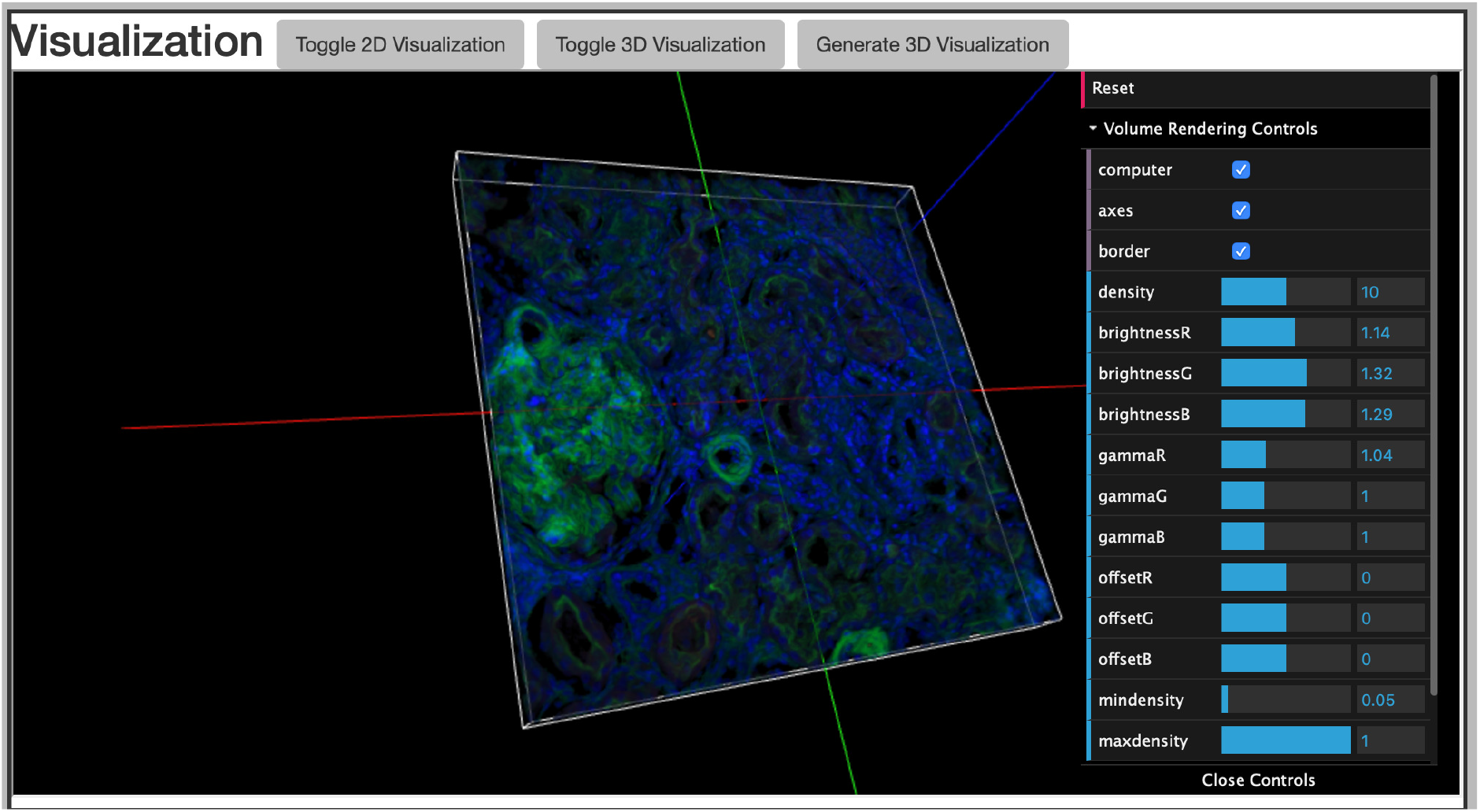
Example of a 3D Visualization. Users are able to adjust the brightness, gamma, and offset of each of the displayed channels in RGB. Users can also zoom in and rotate the volume.

**Figure 4.**
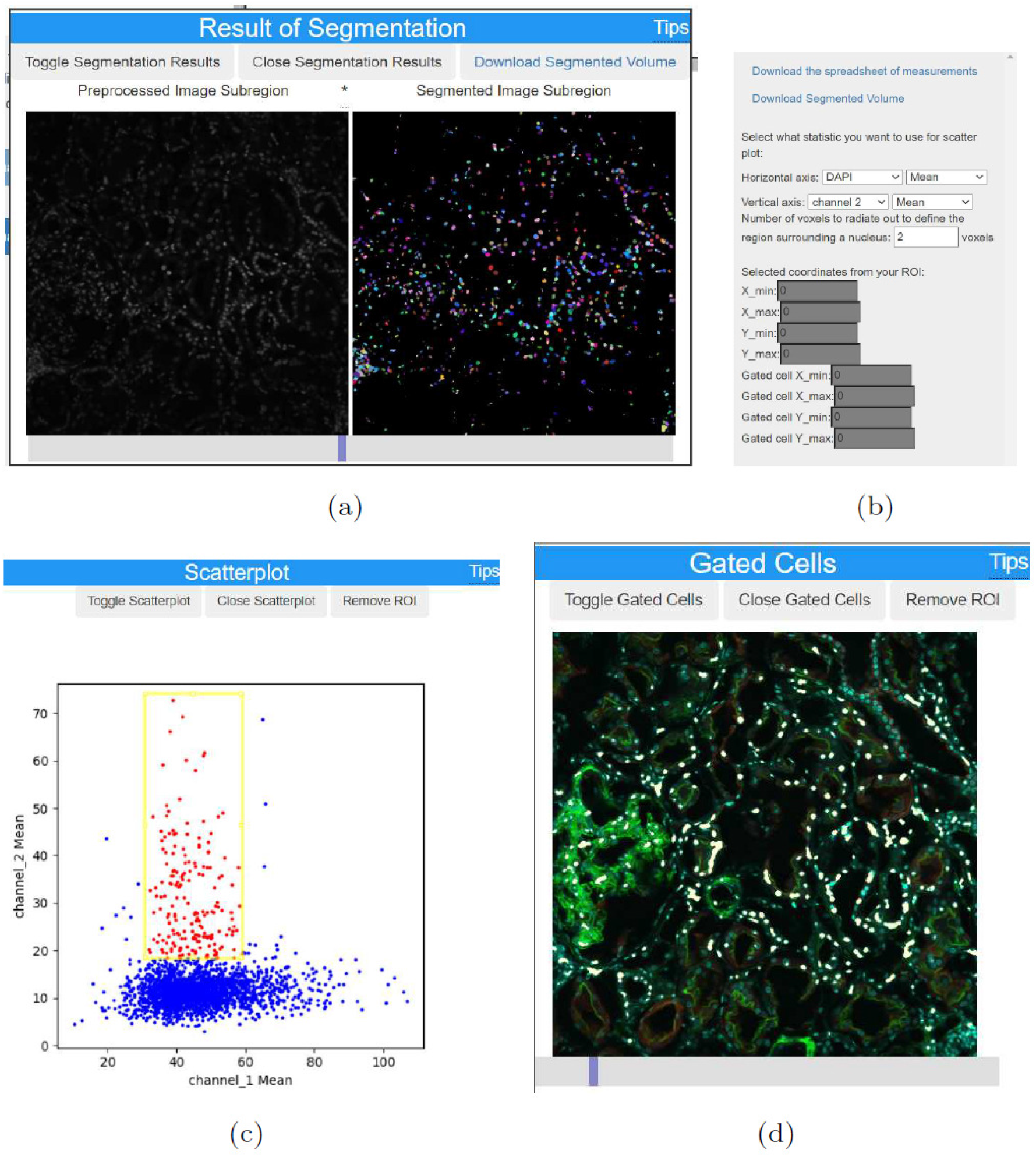
Examples of quantitative image analysis in DINAVID with scatter-plot and plotting gated nuclei (a) Example segmentation using watershed [33], (b) Panel for quantitative scatter plot settings, (c) Scatter plot of nuclei, (d) Mapping of gated nuclei.

## Conclusion

In this paper, we described the DINAVID system that was designed and developed, using open source tools, for the analysis and visualization of 3D microscopy volumes. The goal is to provide a system capable of analyzing large 3D microscopy volumes using sophisticated machine learning methods that biologists can use without worrying about managing computational resources. The results can also be visualized in 2D or 3D. We also compared DINAVID to existing systems and feel that the approaches we have taken eases the burden on users and provides them with an extensible set of tools for 3D volume analysis. The source code and user access to DINAVID are available upon request.

In the future we will deploy more image analysis tools on DINAVID including additional machine learning architectures for microscopy image analysis. One example that can be added in the future is the ability to provide a way for a user to upload training data into DINAVID so that the machine learning models can be retrained. We also plan to add features to DINAVID to process more types of microscopy image data, such as cell classification, transformer based segmentation, and supporting additional bio-imaging formats beyond 3D composite TIFFs or collections of 2D slices. We are also investigating adding online transfer learning tools that will allow users to investigate their own types of data by training machine learning methods.

## Appendix

### Appendix A: DINAVID System: The Details Inside

One can request access to the DINAVID system by sending an email to. The DINAVID source code package is also available upon request to. The source code package includes installation scripts and a list of the appropriate dependencies. Several tutorials are also included along with a complete description of how the DINAVID system works.

DINAVID supports the integration of new image analysis methods for both pre-processing and segmentation. Note if a machine learning tool is to be added, DINAVID does not support training techniques and it is assumed that a pre-trained model (inference model) is to be added. New image analysis methods can be added to DINAVID as long as they are implemented in Python and properly modified so that the input and output directories are consistent with DINAVID. Any inputs or parameters that are user-provided need to be added as an option to the user interface. The source code package has complete instructions for adding new functions to DINAVID.

The hardware that we have for the current version of DINAVID is as follows: Hardware Configuration

- CPU: Intel Core i7-6900K
- RAM: 128GB
- GPU: NVIDIA Titan XP 12GB RAM per GPU (4 GPUs)
- Storage Driver: 1TB SSD + 10TB HDD

Note that DINAVID system can also be used with a cheaper GPU card for the image analysis.

### Appendix B: Description of Available Pre-Processing Methods

Here we describe the existing pre-processing methods available in the current version of DINAVID. For Gaussian filtering, used to smooth or blur an image, we allow the user to adjust the filter kernel size using the parameter *σ* that controls the width of Gaussian kernel. The median filter can also be used to remove noise from the image by adjusting the window size of the kernel. Median background subtraction is used to remove the background from an image by subtracting a median filtered image from the original image. Thresholding is also available including simple thresholding and Otsu’s method [35]. Clamping removes low intensity values in the image. Pixels whose intensity is lower than the threshold are set to 0, while pixels whose intensity is greater than the threshold remains the same.

Two dimensional morphological operations for binary images, such as erosion, dilation, opening, and closing, are available to determine how local content of the image is shaped relative to a given flat structuring element [36]. The structuring element is defined by its shape and size. In addition, rolling ball [37] is one of the options made available for background subtraction. We use a Python version implemented by [38], wherein users can adjust the radius of the ball used for background subtraction. All functions, with the exception of rolling ball subtraction, are implemented via OpenCV [39].

## Availability and requirements

**Project name:** DINAVID

**Project home page:** https://engineering.purdue.edu/~micros/

**DINAVID home page:** https://photon.ecn.purdue.edu/~micros/

To request an account to use the DINAVID system please send an email to.

**Source Code:** The DINAVID source code is available upon request to

**Operating system:** Linux

**Programming language:** Python, Java, JavaScript, HTML, CSS

**Other requirements:** Python Libraries Required: amqp, asgiref, asn1crypto, billiard, celery, certifi, cffi, chardet, click, click-didyoumean, click-repl, cryptography, cycler, Cython, dataclasses, decorator, Django, django-jquery, funcsigs, future, h5py, idna, imageio, importlib-metadata, iniconfig, intel-openmp, kiwisolver, kombu, ldap3, matplotlib, mkl, mod-wsgi, networkx, numpy, opencv-python, Pillow, pip, prompt-toolkit, pyasn1, pycparser, pyparsing, python-dateutil, pytz, PyWavelets, scikit-image, scipy, setuptools, six, sqlparse, tbb, tifffile, torch, torchfile, torchvision, typing-extensions, vine, wcwidth, wheel, zipp

**License:** The source code is distributed under Creative Commons license Attribution-NonCommercial-ShareAlike - CC BY-NC-SA

The source code is available on request to. The source code package includes installation scripts and list of the appropriate dependencies.

**Any restrictions to use by non-academics:** You may not use the source code for commercial purposes as determined by Creative Commons licenses CC BY-NC-SA as indicated above.

## Abbreviations

DINAVID: Distributed and Networked Analysis of Volumetric Image Data
3D: Three Dimensional
2D: Two Dimensional
MB: Megabyte
GB: Gigabyte
TB: Terabyte
GPU: Graphics Processing Unit
WIPP: Web Image Processing Pipeline
NIST: National Institute of Standards and Technology
CNN: Convolutional Neural Network
WebGL: Web Graphics Library
Mb/s: Megabit per Second
HTML: Hypertext Markup Language
CSS: Cascading Style Sheets
AJAX: Asynchronous JavaScript and XML
ROI: Region of Interest
GUI: Graphical User Interface;

## Acknowledgements

Tissue was stained and imaged at the Indiana Center for Biological Microscopy as described previously in [40].

## Authors’ contributions

Project Conceptualization: EJD KWD PS. Project Funding acquisition: EJD KWD PS. Project Administration and Supervision: EJD. System Development: SH AC SL CF CY LW EJD. Testing, Performance analysis: SH AC SL CF CY LW. Testing, User-experience: KWD SW TMA. Writing—original draft: SH AC SL CY LW EJD PS. Writing—review and editing: SH AC LW SW KD PS EJD. All authors read and approved the final manuscript.

## Availability of data and materials

The DINAVID source code is available upon request to. You may not use the source code for commercial purposes as determined by Creative Commons licenses CC BY-NC-SA. The image volume used in this paper is available at [41].

## Declarations

### Ethics approval and consent to participate

Human tissue was collected and processed under the Institutional Review Board at Indiana University approved protocol 1906572234.

## Authors’ information

Shuo Han is a former PhD Student in Electrical and Computer Engineering at Purdue in West Lafayette, Indiana. Her research interests include image processing, microscopy image analysis, computer vision, and machine learning.

Alain Chen is a PhD Student in Electrical and Computer Engineering at Purdue in West Lafayette, Indiana. His research interests include image processing, microscopy image analysis, computer vision, and machine learning.

Soonam Lee is a former PhD Student in Electrical and Computer Engineering at Purdue in West Lafayette, Indiana. His research interests include image processing, microscopy image analysis, computer vision, and machine learning.

Chichen Fu is a former PhD Student in Electrical and Computer Engineering at Purdue in West Lafayette, Indiana. His research interests include image processing, microscopy image analysis, computer vision, and machine learning.

Changye Yang is a PhD Student in Electrical and Computer Engineering at Purdue in West Lafayette, Indiana. His research interests include image processing, microscopy image analysis, computer vision, and machine learning.

Liming Wu is a PhD Student in Electrical and Computer Engineering at Purdue in West Lafayette, Indiana. His research interests include image and video processing, microscopy image analysis, computer vision, and machine learning.

Kenneth Dunn is a Professor of Medicine and Biochemistry at the Indiana University School of Medicine. His research interests include cell biology, animal physiology and methods of optical microscopy and digital image analysis.

Seth Winfree is a Post-doctoral Research Associate in Pathology and Microbiology at University of Nebraska Medical Center in Omaha, Nebraska. His research interests include precision medicine, kidney immunology, kidney disease, tissue cytometry/histocytometry, transcriptomics, microscopy image analysis, machine learning and multimodal data integration.

Tarek M. El-Achkar is the David M. and Julie B. DeWitt Professor of Nephrology Research and Professor of Medicine at Indiana University School of Medicine in Indianapolis, Indiana. His research interests include precision medicine, Uromodulin biology, kidney immunology, kidney disease, tissue cytometry/histocytometry, transcriptomics, microscopy image analysis, and machine learning.

Paul Salama is a Professor of Electrical and Computer Engineering as well as the Associate Dean for Graduate Programs in the Purdue School of Engineering and Technology at Indiana University-Purdue University, Indianapolis. His research interests are in Signal/Image Processing, Biomedical Image Analysis, Machine/Deep Learning, Image/Video Compression, and the Reliable/Secure Transmission of Compressed Data.

Edward J. Delp is the The Charles William Harrison Distinguished Professor of Electrical and Computer Engineering and Professor of Biomedical Engineering at Purdue University in West Lafayette, Indiana.

His research interests include image and video processing, image analysis, computer vision, machine learning, image and video compression, multimedia security, medical imaging, multimedia systems, communication and information theory.

